# Bias and measurement error in comparative analyses: a case study with the Ornstein Uhlenbeck model

**DOI:** 10.1101/004036

**Authors:** Gavin H. Thomas, Natalie Cooper, Chris Venditti, Andrew Meade, Rob P. Freckleton

**Affiliations:** Department of Animal and Plant Sciences, University of Sheffield, Sheffield S10 2TN, UK; School of Natural Sciences, Trinity College Dublin, Dublin 2, Ireland and Trinity Centre for Biodiversity Research, Trinity College Dublin, Dublin 2, Ireland; School of Biological Sciences, University of Reading, Reading, Berkshire, RG6 6BX, UK

## Abstract

Phylogenetic comparative methods are increasingly used to give new insight into variation, causes and consequences of trait variation among species. The foundation of these methods is a suite of models that attempt to capture evolutionary patterns by extending the Brownian constant variance model. However, the parameters of these models have been hypothesised to be biased and only asymptotically behave in a statistically predictable way as datasets become large. This does not seem to be widely appreciated. We show that a commonly used model in evolutionary biology (the Ornstein-Uhlenbeck model) is biased over a wide range of conditions. Many studies fitting this model use datasets that are small and prone to substantial biases. Our results suggest that simulating fitted models and comparing with empirical results is critical when fitting OU and other extensions of the Brownian model.

## Introduction

The outcomes of evolutionary processes acting over millions of years can be seen in the distributions and covariances of species traits at the tips of phylogenetic trees. Such processes can be modeled using phylogenetic comparative methods (PCMs). These approaches have been used, for example, to model niche conservatism (Wiens et al. 2010), infer potential rates of species responses to climate change (Quintero and Wiens 2013), test the role of ecological niche as a driver of morphological evolution (Pienaar et al. 2013), compare rates of evolution of species traits (Claramunt et al. 2012) and test for constraints in adaptive radiations (Blackburn et al. 2013). PCMs are now widely recognized as powerful tools with which to model observed patterns in species traits and potentially to infer the evolutionary processes that drive them (e.g., Freckleton 2009; Nunn 2011; O'Meara 2012; Pennell and Harmon 2013). Despite these advances, the properties of some of the most widely used PCMs are poorly understood leading to the potential for inappropriate use and misinterpretation of results.

The majority of PCMs use an explicit evolutionary model to characterize trait evolution (Freckleton et al. 2011). A model can be fit using statistical methods (typically maximum likelihood) and model parameters interrogated to test ideas from evolutionary theory. Most model-based methods for characterizing trait evolution are based on the Brownian constant variance model (for exceptions see Price et al. 1997; Harvey and Rambaut 2000; Freckleton and Harvey 2006). The Brownian model, first applied in a phylogenetic context by Cavalli-Sforza and Edwards (1967) and to across-species data by Felsenstein (1973), is a simple model of trait evolution in which trait variance accrues as a linear function of time, and makes the prediction that traits of closely-related species are more similar than those of distantly-related ones. The Brownian model has been modified in various ways to account for a suite of ecological and evolutionary processes (e.g., Grafen 1989; Hansen 1997; Pagel 1997, Pagel 1999). Most of these involve a transformation of the tree and thereby fitting a model with one or more extra parameters. These modified Brownian models have become very popular because they fit better, have attractive biological interpretations, and are implemented in freely available packages (e.g., R; R Development Core Team 2013).

The modification of Brownian models to account for non-Brownian processes can be traced back to Grafen (1989) who introduced a parameter *ρ* to account for non-linear scaling of evolutionary changes. *ρ* is a power transformation of node heights in the phylogeny, and is fitted by maximum likelihood. It is worth noting that Grafen warned explicitly about potential drawbacks with *ρ*. Specifically: (i) the parameter is likely to be biased; (ii) its asymptotic sampling variance is non-zero; and (iii) values of the parameter should be interpreted cautiously. Freckleton *et al*. (2002) showed that *ρ* was indeed biased as predicted by Grafen (1989). They also showed that another parameter, Pagel’s λ (Pagel 1997, Pagel 1999), also exhibited biases (but that these were likely to have minimal consequences in analyses using reasonable sample sizes). This strongly suggests that simulations are required to test the properties of such parameters, and that as recommended by Grafen (1989) “careless” parameterization of the phylogenetic variance-covariance matrix should be avoided.

One of the most commonly used Brownian-like models is the Ornstein Uhlenbeck (OU) model. This is a modification of the Brownian model with an additional parameter *α* that measures the strength of return towards a theoretical optimum (Hansen 1997). The popularity of the OU model has grown exponentially in recent years (Fig. 1), in part because these models are now easy to implement via packages in R (e.g. ouch, GEIGER and OUwie; Butler and King 2004; Harmon et al. 2008; Beaulieu and O'Meara 2012). OU models have become particularly prevalent in ecological studies: 652 papers containing the phrases “Ornstein Uhlenbeck” and “ecology” were published between 2011 and 2013 (Google Scholar search 30^th^ January 2014). Additionally, although they are pattern based, fit to an OU model is now being used as evidence for the action of ecological, or ecologically driven, processes such as phylogenetic niche conservatism, convergent evolution and stabilizing selection (e.g., Wiens et al. 2010; Christin et al. 2013; Ingram and Mahler 2013).

**Figure 1.**
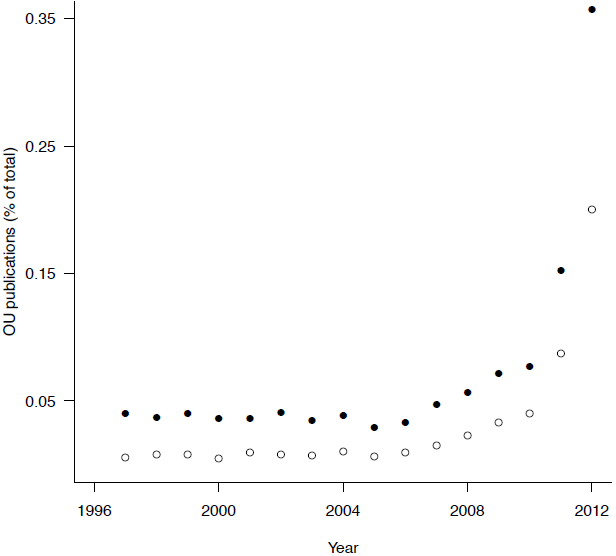
*The number of evolutionary biology (closed circles) and ecology (open circles) papers published between 2005 and 2012 containing the phrase “Ornstein-Uhlenbeck”, as a proportion of the total number of evolutionary biology or ecology papers published that year*.

Here we investigate the OU model because it does not appear to have been subject to a rigorous critique. The *α* parameter is very similar to Grafen’s *ρ* (along with other parameters in the literature) in that it is in effect a non-linear transformation of the phylogeny. Consequently, one might imagine that it shares some of the same statistical properties. Results from Ho & Ané (2013), indicate that there is the potential for biases in *α*, but the implications for practice (e.g. likelihood of rejection of alternative models) is unexplored other than to note that tree shape may play an influence. We present simulations demonstrating the inherent bias in estimating the *α* parameter, discuss the intricacies of interpreting OU models biologically, and provide advice for appropriate use of OU models in comparative analyses. We also show that very small amounts of measurement error in data can have profound effects on the performance of models. Many of our findings are applicable to other models of evolution (see Discussion and Supplemental Figs. S6– S11), but we focus here on the OU model because of the recent increase in publications using the model and because of the underappreciated ambiguity in the link between pattern and process when interpreting estimates of the *α* parameter.

## Materials and Methods

### Model Outline

According to the Brownian model, a trait X evolves at random at a rate *σ*

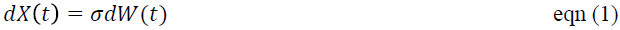

where *W(t)* is a white noise function and is a random variate drawn from a normal distribution with mean 0 and variance *σ^2^*. This model assumes that there is no overall drift in the direction of evolution (hence the expectation of *W(t)* is zero) and that the rate of evolution is constant. The model has two parameters, *σ* and the state of the root at time zero, *X*(0). The Brownian model predicts after a time *T*the variance in trait value *X_i_* for species *i* is.

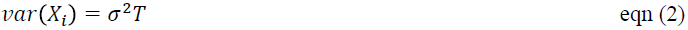

and the covariance in traits for species *i* and *j* is:

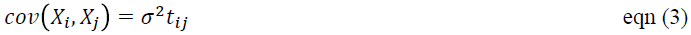

where *t_ij_* is the shared evolutionary pathway for species *i* and *j*, i.e. the time at which they last shared a common ancestor. Equations (2) and (3) encapsulate the simplicity of the Brownian model, namely it predicts that variances accrue as a linear function of time.

The Ornstein-Uhlenbeck (OU) model describes a mean-reverting process and has the following form, adding an extra term to the Brownian model:

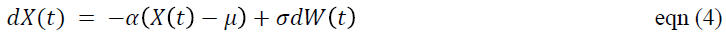

The parameter *μ* is a long-term mean, and it is assumed that species evolve around this value. *α* is the strength of evolutionary force that returns traits back towards the mean if they evolve away. The OU model was introduced to population genetics by Lande (1976) to model stabilizing selection in which the mean was recast as a fitness optimum on an adaptive landscape. The process operating in comparative data is analogous, although clearly is not stabilizing selection despite being sometimes referred to as such (see Discussion). This model has two parameters in addition to those of the Brownian model, *α* and *μ*.

The OU model predicts that after a time *T*for a species *i*, the variance in trait value *X_i_* is:

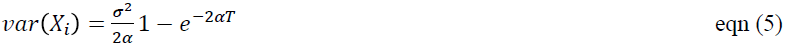

And for a pair of species *i* and *j*, the covariance in traits is:

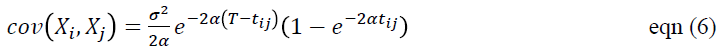

The variances and covariances predicted by equations (5) and (6) are more complex than those predicted by the Brownian model. In the light of the results below, some properties of this model are worth highlighting:

i. *If α is small then evolution is approximately Brownian*: If *α* is small then 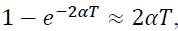, i.e. traits accrue variance as if evolving according to a Brownian process. The question is obviously what is a small value of *α*? Figure 2 provides some guidance on this issue, and see point (iv) below.
ii. *If species i and j diverged recently, evolution is approximately Brownian:* if two species diverged recently, then *T — t_ij_* ≈ 0 and hence 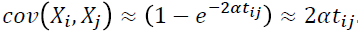 Thus, recently diverged species provide little information relevant to estimating non-Brownian evolution according to an OU process.
iii. *In the long-term the imprint of history is weakened:* if *T* is large (i.e. evolution proceeds for a long time), equation (3) predicts that the variance in *X_i_* tends to a constant, i.e. because the expected value of *X_i_* is *μ*. Similarly, in equation (4), the covariance between traits tends to a constant because *T* becomes large relative to *t_ij_*. Consequently for large groups the model implies that the imprint of history is weak. As a result adding more species to a phylogeny will contribute only relatively weakly to fitting the model, as the deeper branches within a phylogeny will contribute little information to estimating the parameters.

Figure 2 gives some numerical examples, chosen to outline the behavior of *α* in relation to the above. We show the predicted covariance in traits (equation 6) scaled by the overall variance (equation 5) plotted against the time since divergence, *t_ij_* scaled by tree height. Figure 2 highlights that the behavior of *α* can be classified into three regions. When *α* is low the model behaves very much like Brownian motion, with the covariance scaling approximately linearly with time. Only as the value becomes moderately larger (in this case approaching a value of 0.5), does the non-Brownian behavior become apparent. Secondly, there is a region of moderate values in which the model leads to markedly non-Brownian patterns of trait variation (Fig. 2B). Finally, when α is large the model behaves differently again: in this case, all imprint of history is lost and the evolution is essentially a rapid burst at the present (Fig. 2C). This will be indistinguishable from phylogenetically unstructured data.

**Figure 2.**
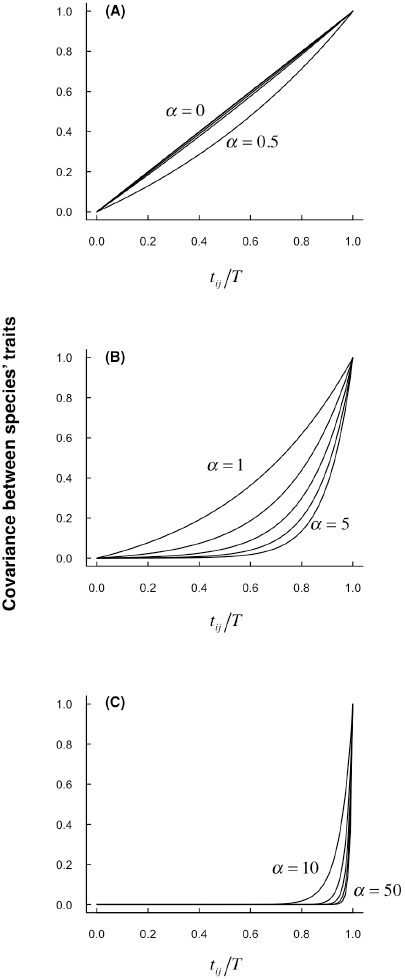
B*Scaling of expected trait similarity with time since evolutionary divergence predicted by the Ornstein-Uhlenbeck model. The covariance between species’ traits values is scaled by the intra-specific trait variance (i.e. equal to correlation between species' traits). This is plotted against the relative time of shared history (time at which species branched from each other, divided by the total tree height).*

In terms of statistical model fitting, a feature to highlight is that in Figure 2A and Figure 2C, changing the value of *α*across large *relative* ranges at the extremes (0 – 0.5 in Fig. 2A; 10-50 in Fig. 2C) produces relatively little quantitative change in the model behavior, and it is in the middle region of parameter space (1 – 5) that the biggest discernable effects occur.

The point of highlighting these aspects of expected behavior is that we believe these properties need to be understood if we are to build up a full picture of when the model is likely to be well estimated, as well as to understand the bounds of what can be inferred from real data.

### Simulations

We used simulations to explore the statistical properties of *α*. We simulated phylogenies with 25, 50, 100, 150, 200, 500, or 1000 tips under pure birth, constant-rate Birth-Death (extinction fractions of 0.25, 0.5 and 0.75), or temporally varying speciation rate (speciation rate modeled as time from the root raised to the power 0.2, 0.5, 2 and 5) models. We simulated 1000 phylogenies for each combination of tips and models resulting in 56,000 simulated phylogenies in total. Trees were simulated using the R package TESS (Hoehna 2013). We then simulated the evolution of a single trait under a Brownian motion model on each phylogeny using the R package MOTMOT (Thomas and Freckleton 2011). We estimated α and compared the fit of a Brownian model to that of an OU model using: (i) a likelihood ratio test with 1 degree of freedom with the transformPhylo.ML function in MOTMOT; and, (ii) with Bayes factors estimated from a stepping stone sampling procedure (Xie et al. 2011) implemented in BayesTraits. The marginal likelihoods of the models were calculated using a stepping stone sampler in which fifty stones were drawn from a beta distribution with (alpha = 0.4 and beta = 1). Each stone was sampled for 20000 iterations (with the first 5000 iterations discarded). We treated Bayes factors > 2 as evidence favouring the OU model. We used three alternative sets of priors on *α*: (i) an exponential distribution with mean = 1; (ii), an exponential distribution with mean = 10, and (iii) a uniform distribution bounded at 0 and 20. For all analyses we used a uniform -100 to 100 prior for *μ* and uniform 0 to 100 for *σ*^2^. We also normalized the tree length to prevent the *α* parameter of the OU model from interacting with *σ*^2^.

OU models have not previously been widely applied within a Bayesian framework, although some packages offer this functionality (e.g. diversitree; FitzJohn 2012) and the choice of prior has not been fully explored. We present the results from the exponential prior with mean = 10 in the main text since this is a broad, liberal prior. We provide results derived from a strong exponential prior with mean = 1 and a bounded uniform prior (range 0–20) as supplementary material (Figs. S2–S5). We ran the MCMC chains for 1×10^6^ iterations, disregarding the first 1×10^4^ as burn-in. Following burn-in the chains were sampled every 1000 to ensure independence of each consecutive sample. Multiple independent chains were run for each analysis to ensure convergence was reached.

We used the same procedure to simulate trait data under a Brownian motion model with known error. Specifically, we simulated trees under a Yule model with 25, 50, 100, 150, 200, 500, or 1000 tips and added branch length of (i) 1%, (ii) 5% or (iii) 10% of the tree height to the tips of the simulated trees. We then simulated data under a Brownian model on each tree. We compared the fit of a Brownian model to that of an OU model (with ML and MCMC stepping stone sampling as above) using the original trees without the addition of extra branch length to the tip edges. The expectation is that the OU model should fit the data better than the Brownian model but the reason for the better fit would be entirely unrelated to any evolutionary process.

Although our focus is on the OU model we also explored three other commonly used models (*κ*, *λ*, and *δ*; Pagel 1997, Pagel 1999) fitted using maximum likelihood using the same simulated data. These models are designed to test for speciational evolution (*κ*), strength of phylogenetic signal (*λ*) and accelerating or decelerating evolution (*δ*). The point of these additional tests is to assess the extent to which parameter behaviors vary across models and the results are reported in the supplementary material.

### Literature Review

To get an overview of the use of OU models in ecology and evolution, we used Google Scholar (accessed 21^st^ November 2013) to locate papers published between 2005 (when the R package ouch was released; Butler and King 2004) and 2013 that contained the terms “Ornstein Uhlenbeck” and either “ecology”, or “evolution” and “biology” (the “biology” term was added to omit physics papers which also use the term “evolution”). We also recorded the total number of papers containing the terms “ecology”, or “evolution” and “biology” published between 2005 and 2013 and plotted the number of OU papers published each year as a proportion of the total number of papers published (Fig. 1).

Next we filtered our Google Scholar search results to focus on empirical papers using OU models (rather than methods papers) published in the following ecology and evolution journals: The American Naturalist, Ecology Letters, Evolution, Journal of Evolutionary Biology, Nature, Proceedings of the National Academy of Sciences, USA, Proceedings of the Royal Society B: Biological Sciences, and Science. For each of these papers we recorded the number of species in the analysis, the study group (amphibians, birds, fish, mammals, reptiles, invertebrates or plants), the statistical package or specific R package used to fit the models, and the reason the authors state for using an OU model (ancestral state reconstructions, detecting convergent evolution, controlling for phylogeny, selecting a model of trait evolution, or other). Where papers included multiple analyses using different numbers of species we used the median number of taxa. Where papers had multiple study groups, statistics/R packages or reasons for fitting OU models we counted them in each relevant category. We summarise these results in Figure S1 and the full dataset is available in Supplemental Table S1 along with the full list of references.

## Results

### Simulations

Figure 3 shows some examples of profile likelihoods for *α* generated on different datasets that were simulated according to a Brownian model. It is clear that the shape of the tree affects the shape of the likelihood profile. Trees with increasing speciation rates from root to tip (i.e., that have a lot of recently evolved species) generate very wide, flat likelihood profiles (Fig. 3A). On the other hand, trees with declining speciation rates from root to tip (Fig. 3B), or generated according to a Yule process (Fig. 3C), yield more strongly peaked profiles.

**Figure 3.**
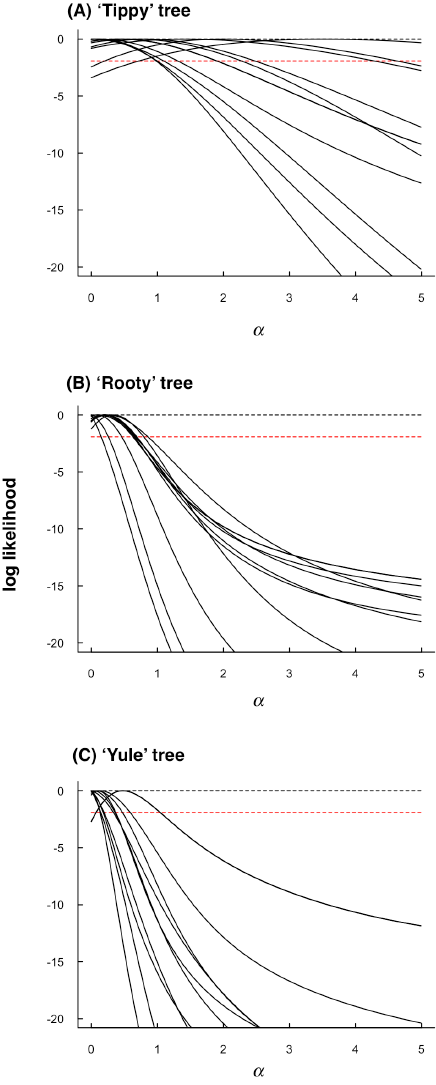
*Examples of profile likelihoods for selected simulated datasets. In all cases the 'true’ value of alpha is 0.*

In all three cases, there are several features worth noting. First, the profiles are asymmetric. There is a hard bound at *α* = 0, and as *α* becomes large, the rate of decrease in the likelihood slows. This is because, as shown in Figure 2, as values of *α* become large, the effects of changing *α* on model predictions are increasingly small. It is also notable, that the Maximum Likelihood estimates of *α* are, in most cases, between 0 and 1. In Figure 2 it was noted that in this region of parameter space, the OU model is very difficult to distinguish from Brownian. This, combined with the asymmetric nature of the profile likelihood is suggestive that estimates of *α* are likely to be biased, and that the asymptotic assumptions of likelihood ratio tests on *α* are likely to be violated.

The effects of the asymmetric profile likelihoods become clear when calculating Type I error for the OU model based on ML estimates across a range of tree shapes and sizes (Fig. 4; see Supplemental Figs. S2 and S3 for results with alternative priors for the Bayesian analyses). Regardless of tree shape, Type I error rates are unacceptably high when tree size is small (Figs. 4A, 4C and 4E). For some tree shapes (particularly where speciation rates accelerate towards the present), Type I error remains >0.05 even for trees with 1000 tips. Bayesian estimation is not subject to bias in the log-likelihood statistic and consistently rejects the OU model. However, estimates of α are similar regardless of method.

**Figure 4.**
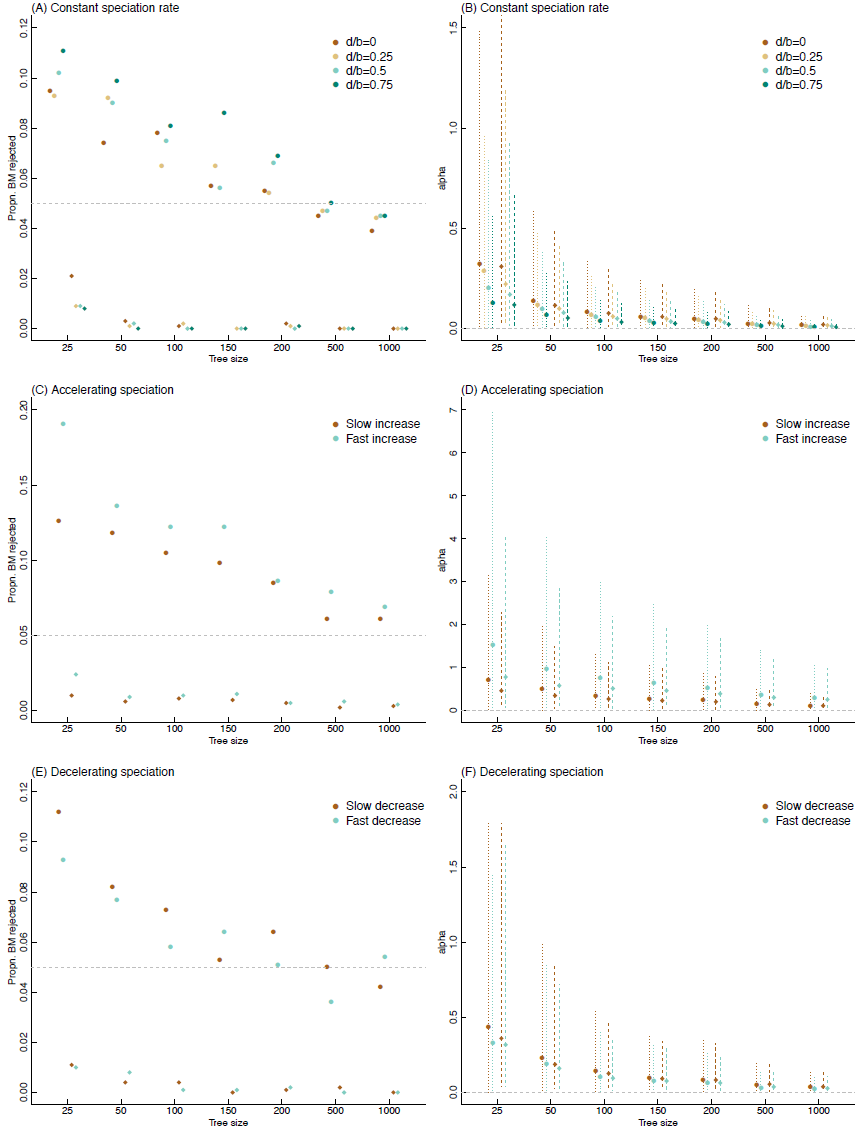
*Rejection rates and estimation of α for different tree sizes and shapes. Rejection rates (a,c,e) for the OU model at multiple tree sizes for models fitted using ML (circles) and Bayesian (diamonds) methods. Estimates of α (b,d,f) based on ML (circles show means and dotted lines show 95% sampling interval from 1000 trees) and Bayesian (diamonds show means and dotted lines show 95% sampling interval of the modal value from the posterior distribution from 1000 trees)*.

Figure 5 shows the proportion of data sets in which the OU model is favoured over the Brownian model for data simulated under Brownian motion with error (also see Figs. S4 and S5). The expectation is that the OU model should fit better because the branch length transformation partially captures the non-Brownian component (the error). There are two points worth noting. First, the frequency with which the OU model is favoured increases with tree size for both ML and Bayesian estimation (Fig. 5A). With as little as 5% error, the OU model becomes extremely difficult to reject, even for trees with just 100 species. This is important for the interpretation of the OU model. We cannot conclude anything about evolutionary process from a single optimum OU model unless error is adequately accounted for. Second, for moderate amounts of error (5-10%), the ML estimates of *α* are consistently >1 (Fig. 5B). Large values of α are similarly difficult to interpret because they are indicative that the signal of the past has been overwritten.

**Figure 5.**
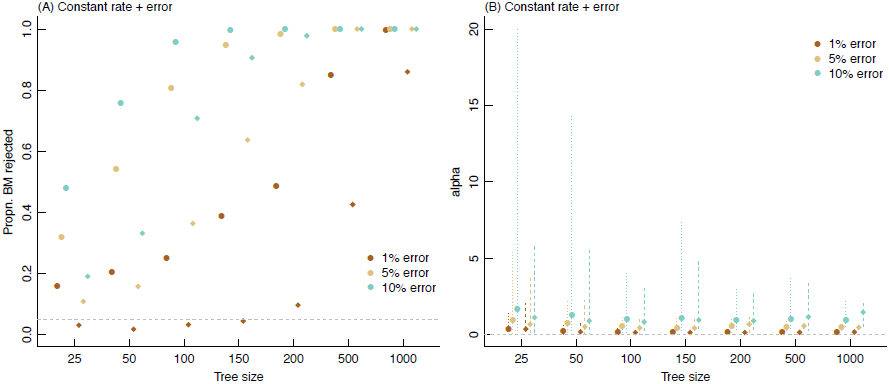
*Rejection of BM with measurement error (a) and estimation of α (b). Symbols follow Figure 4*.

The biases reported above are not unique to the OU model. The parameter *δ*, which tests for accelerating or decelerating rates of evolution (Pagel 1997, Pagel 1999) also suffers from elevated Type I error and is upwardly biased, favoring models that imply temporally increasing evolutionary rates (Supplemental Fig. S8). In contrast, although the speciational model (*κ*) has slightly elevated Type I errors, the parameter estimates are unbiased for the majority of tree sizes and shapes (Supplemental Fig. S6). As previously shown (Freckleton et al. 2002), the *λ* model is conservative with low Type I error rates (Supplemental Fig. S7). As with the OU model, the *δ* models are all strongly favored over Brownian motion when there is error in the species data (Supplemental Figs. S9–S11).

### Literature Review

In total, 3720 papers containing the terms “Ornstein Uhlenbeck” and either “ecology”, or “evolution” and “biology” were published between 2005 and 2013, and the number has increased substantially since 2005 (Fig. 1). Most papers fit OU models to phylogenies with fewer than 100 taxa (mean = 166.97 ± 43.86, median = 58, Supplemental Fig. S1 and (Table S1). The majority of papers fit OU models using R packages, particularly GEIGER and, although other uses are becoming more common, most papers use OU models in an effort to discern the “best” model of trait evolution or to control for phylogenetic non-independence (Supplemental Table S1 and Fig. S1).

## Discussion

Although it is possible to create and implement new models for comparative data that encompass a range of processes, we have to beware that such models are statistically complex and may behave in unexpected ways. Transformations of the variance-covariance matrix in the Brownian model are an attractive and computationally simple way to modify the basic model to include evolutionary processes. But, as first pointed out by Grafen (1989), the statistical consequences of these modifications can include biases and problems with interpretation. The results we have presented here illustrate that such biases can occur under conditions that closely match the size and type of datasets that are commonly used.

In the case of the OU model, we can make a series of specific recommendations for future analyses:

1. *At least 200 species should be included*. The results of the simulations indicate that, in general, analyses based on small datasets are prone to biases that decrease only slowly as the size of the dataset increases.
2. *Likelihood ratio tests are untrustworthy*. We would expect these to be approximate because αis bounded and has a non-linear effect on the expected variances. The simulations indicate that likelihood ratio tests should not be relied upon for analyses with small sample sizes, and that for robust inference and testing, alternatives, such as MCMC, should be considered.
3. *Simulate fitted models and compare these to your empirical results*. A good strategy for data exploration would be to simulate data under BM and fitted OU models to generate distributions of parameters under known values. These can then be compared to results for your dataset (see Slater and Pennell 2013 for a related approach). This is important because we have shown that the shape of a phylogeny has consequences for parameter biases and hypothesis tests. Any given tree will therefore generate unique parameter estimates. Simulating data under the BM model will generate null distributions (e.g., Boettiger et al. 2012). Generating data under the fitted OU model will allow an assessment of whether it is possible to retrieve known values, or whether there is evidence of bias.
4. *Consider plausible alternative hypotheses*. The results indicate that a simple parsimonious model can be mistaken for an OU process: namely a BM process in which a small amount of error is added to the data. Small (<1) values of *α* are quite compatible with this interpretation of the data. The effects of error become more severe with increasing tree size.

In terms of the final point, it would be very useful to have estimates of measurement errors for the species’ observations, however the inclusion of species-specific variances has to be done carefully (e.g., Grafen 1989). It would be tempting to create a vector of variances that could be combined with **V** to create an overall combined variance matrix (O'Meara et al. 2006; Harmon et al. 2010). However, if the variances are large, vary among species or are based on small samples, this can create additional complications for the modeling. This is because the variances will themselves be estimates and subject to error. Treating such variances as error-free can potentially create biases (Freckleton unpub). If the variances are unknown then Pagel’s *λ* (Pagel 1997, Pagel 1999) can be used to estimate the non-Brownian component of the trait variance. However estimating this parameter simultaneously with estimating *α* would have to be undertaken carefully as the two transformations have quite similar effects on the predicted variances.

Our review of the literature revealed that the OU model is frequently described and interpreted as a model of ‘stabilizing selection’. The OU model is attractive because it sets effective bounds in probability on the size of species’ traits, but to use the term ‘stabilizing selection’ is inaccurate and misleading. As formulated by Hansen (1997), a niche has a primary optimum that is the mean of individual species optima for that niche. Under this formulation, α can be considered as the strength of the pull towards the primary optima (Hansen 2012) but not as an estimate of stabilizing selection in the population genetics sense.

In terms of interpretation, it is worth pointing out that data will exhibit few signs of the limits to trait values that result from an OU process. This is because according to the OU process traits move back towards the optimum as their values become particularly large or small. Past history is effectively wiped over, as pointed out above, with the consequence that covariances between species become small (Fig. 2). Indeed, Figure 2 implies that the outcome of the OU process is indistinguishable from an acceleration of the rate of evolution towards the present. This interpretation is particular important for inference of phylogenetic niche conservatism. Although definitions of phylogenetic niche conservatism vary (Losos 2008b, a; Wiens 2008; Crisp and Cook 2012) one proposed approach is to fit an OU model and take rejection of BM in favour of OU as evidence of constraints on niche evolution (Wiens et al. 2010). A more parsimonious explanation would be that there is measurement error in the data. This seems particularly likely for climatic niche traits that are often averaged across an entire species range. Concluding from data on extant species that traits are limited by an OU-like process will therefore always be a leap of faith.

If ancillary data are available, however, this may not be the case. If data on fossils are available then these could be incorporated into the analysis (Slater et al. 2012). A caution here is that the OU model for non-ultrametric trees has to be carefully parameterized because for non-ultrametric trees the co-variances in equation (6) depend on both the shared distances between species and the distance of a node to the nearest tip. This creates potential problems in parameterization and in interpretation because the variance-covariance matrix is no longer treelike (G. Slater unpub.). Some current implementations of the OU model are based on transforming the tree directly, rather than transforming the variance co-variance matrix (e.g., MOTMOT; Thomas and Freckleton 2011). These implementations should not be used with fossil data.

The results we report are not unique to the OU model. As noted above, Grafen (1989) pointed out that his *ρ* parameter would likely be biased, and other transformations (e.g., *λ*, *δ*, ACDC, Pagel 1997, Pagel 1999; Blomberg et al. 2003) will exhibit similar behavior (e.g., see Freckleton et al. 2002). Freckleton *et al.* (2002) explored the behavior of Pagel’s *λ* and found that this statistic was biased but in manner that was likely to be conservative in the rejection of the Brownian model (also see Supplemental Fig. S7). Some transformations may be unbiased: for example, Pagel’s *κ* is a transformation of branch lengths, and is essentially the “standard diagnostic” that is used in testing the assumptions of phylogenetic contrasts, and known to behave appropriately (Supplemental Fig. S6; Garland et al. 1992).

In conclusion, the simulations we report highlight that there are some limits to what we can learn from phylogenies and comparative data on extant species. The OU model is a good example of this: large values of *α* will lead to a loss of history and the dataset will only represent the most recent evolutionary changes. Moreover, different processes can easily yield similar patterns (e.g., Revell et al. 2008) with the consequence that rejection of the Brownian model in favour of another does not necessarily say anything about process. This problem can be alleviated to some extent if model comparisons are set in a firm hypothesis testing framework in which alternative hypotheses make clear predictions of emerging patterns that can be unambiguously associated with particular models (e.g., Cooper et al. 2011). We shouldn’t use any statistical model without thinking carefully about the limits in terms of both data and interpretation. The results presented above highlight that in the case of the OU model, this is especially important.

## Acknowledgements

Thanks to Mark Pagel, Matt Pennell, Graham Slater and Rich FitzJohn for fruitful discussions about OU models. GHT was supported by a Royal Society University Research Fellowship. AM was supported by BBSRC grant BB/K004344/1 and the computing time was funded by European Research Council Grant no. 268744, Mother Tongue.

## Supporting Information 1: Supplemental tables, figures and references

**Table S1.**
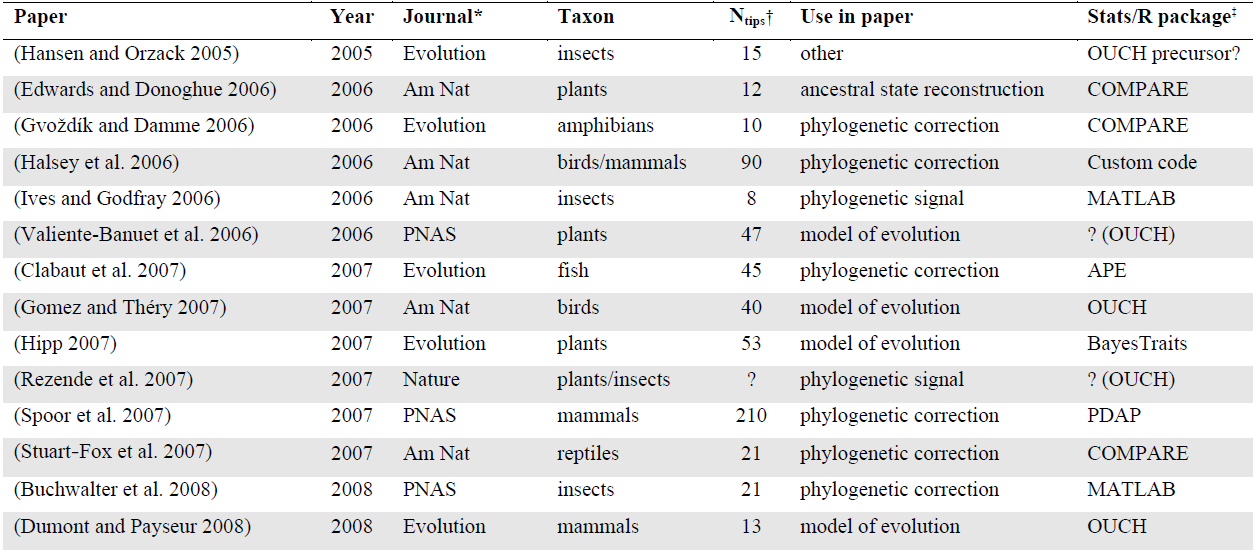
Details of the papers used in our literature review. For each paper we recorded the study taxon, the number of tips in the phylogeny used to fit the Ornstein Uhlenbeck (OU) model, how the authors used the OU model in the paper, and the statistical package (usually R) used to carry out the analyses. For a full reference list see below.

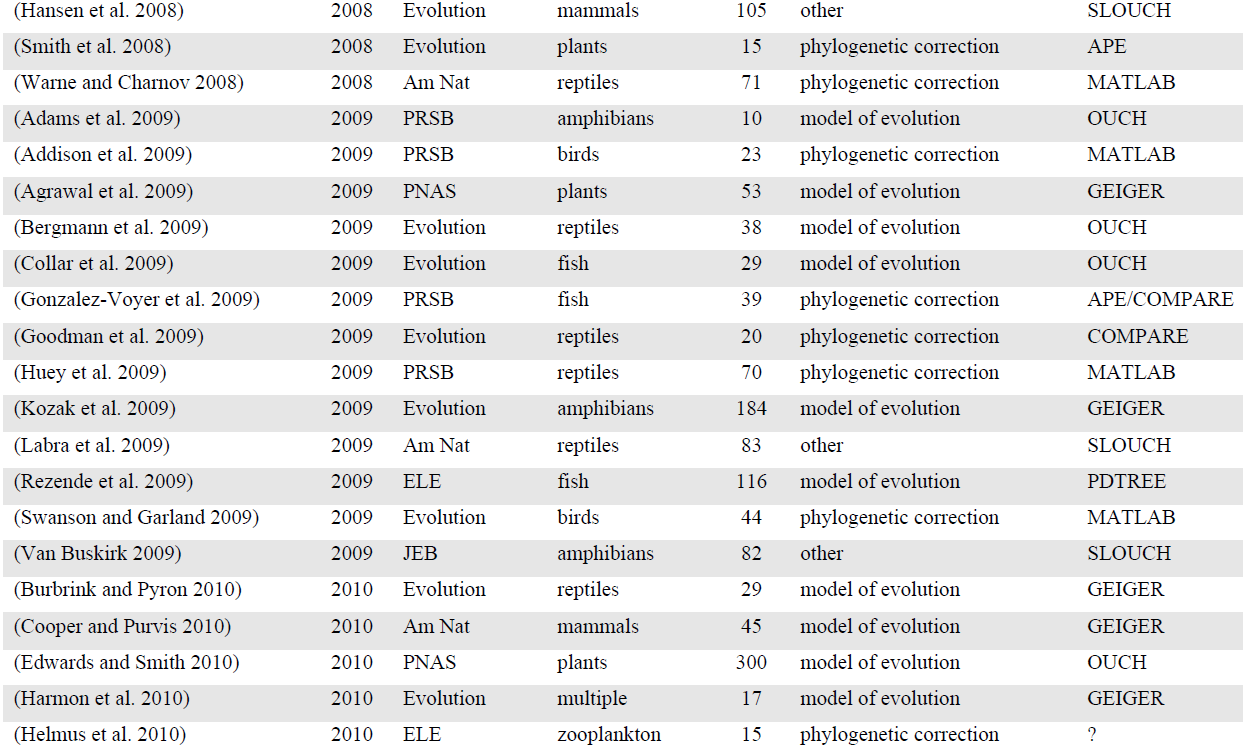

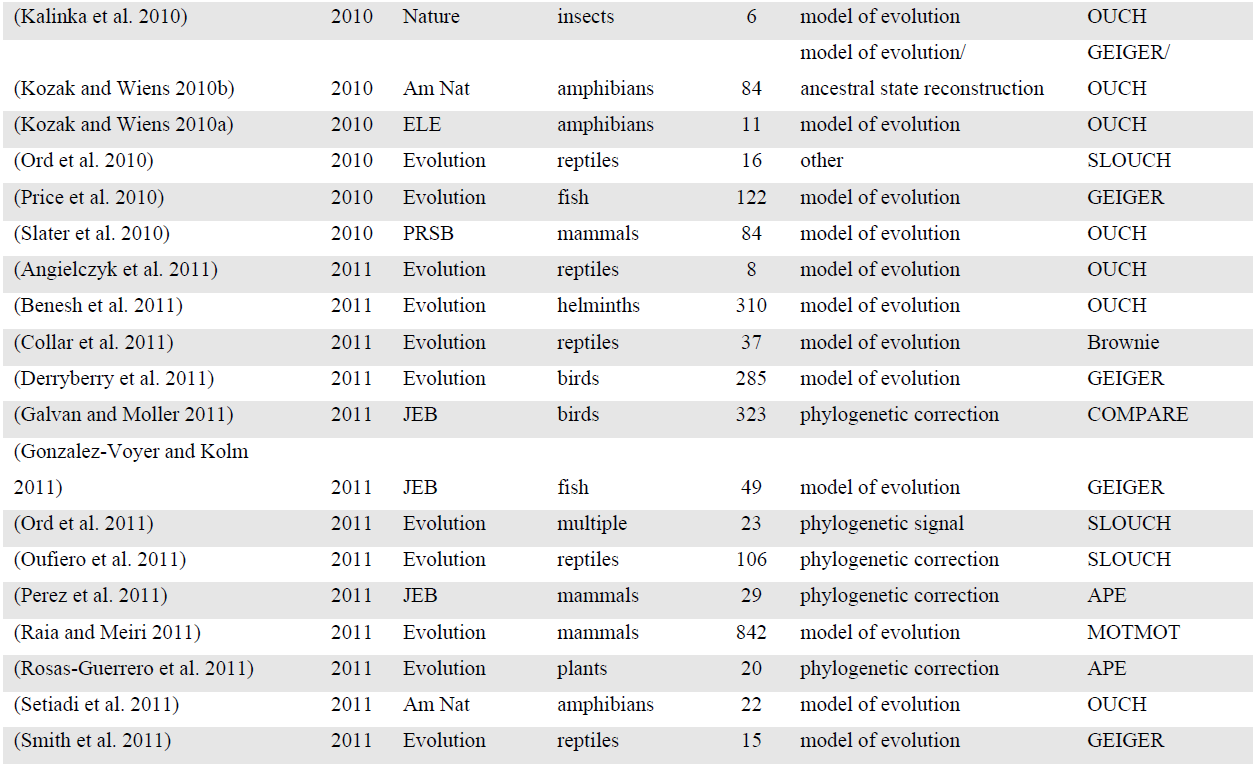

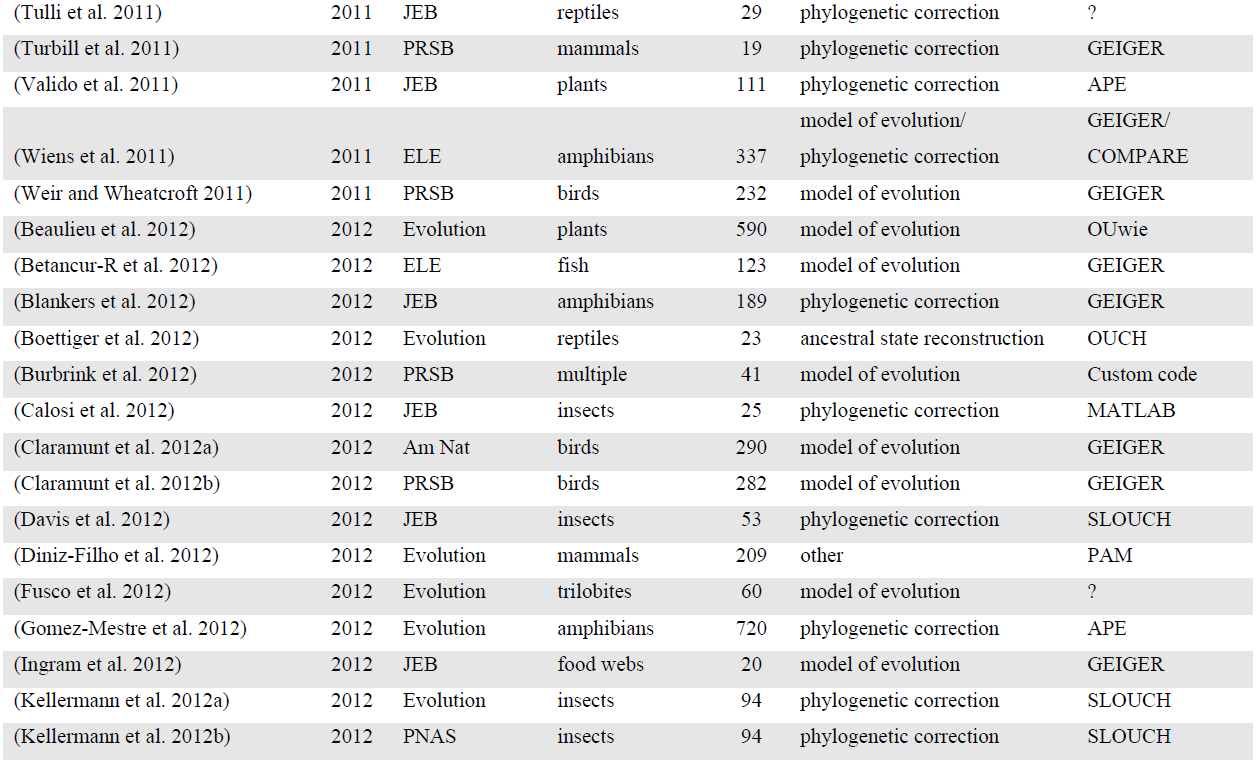

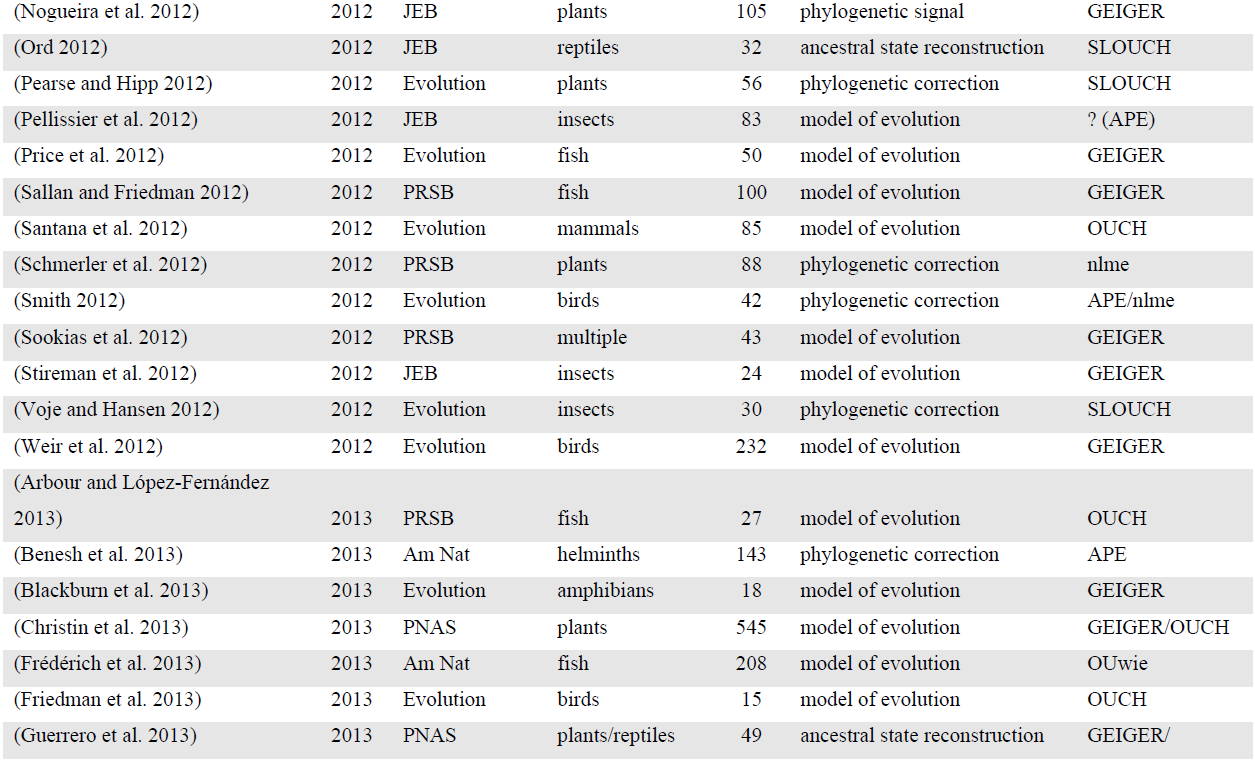

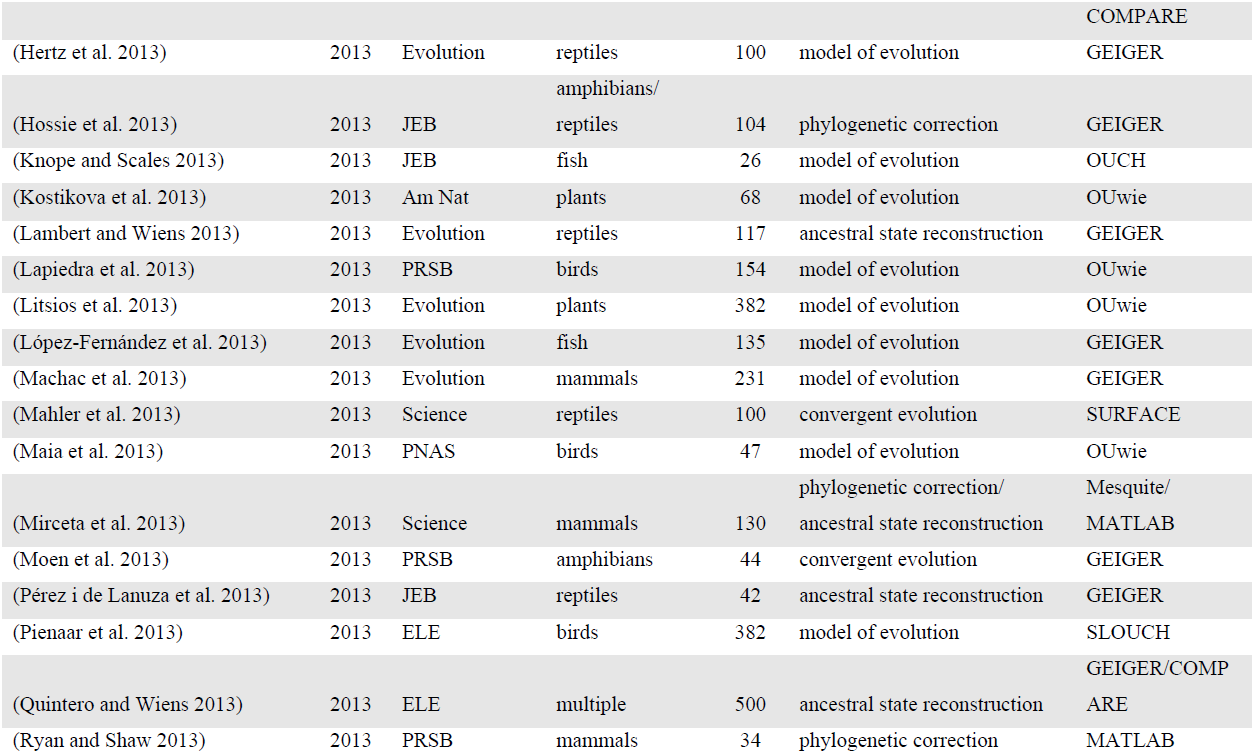

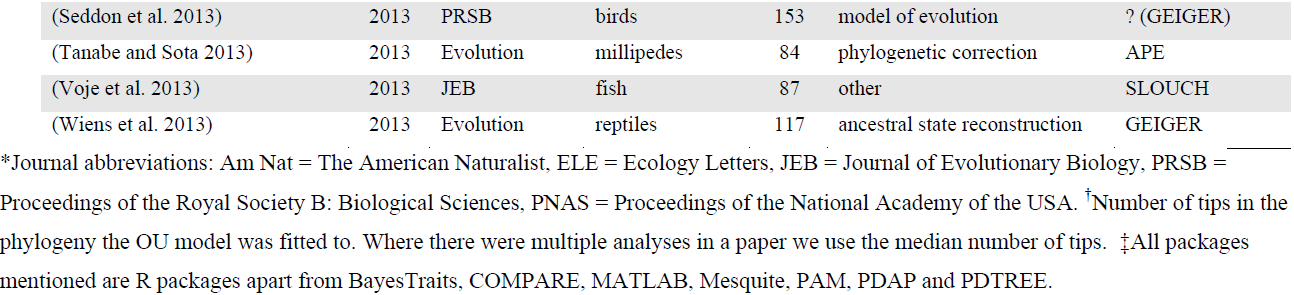

**Figure S1.**
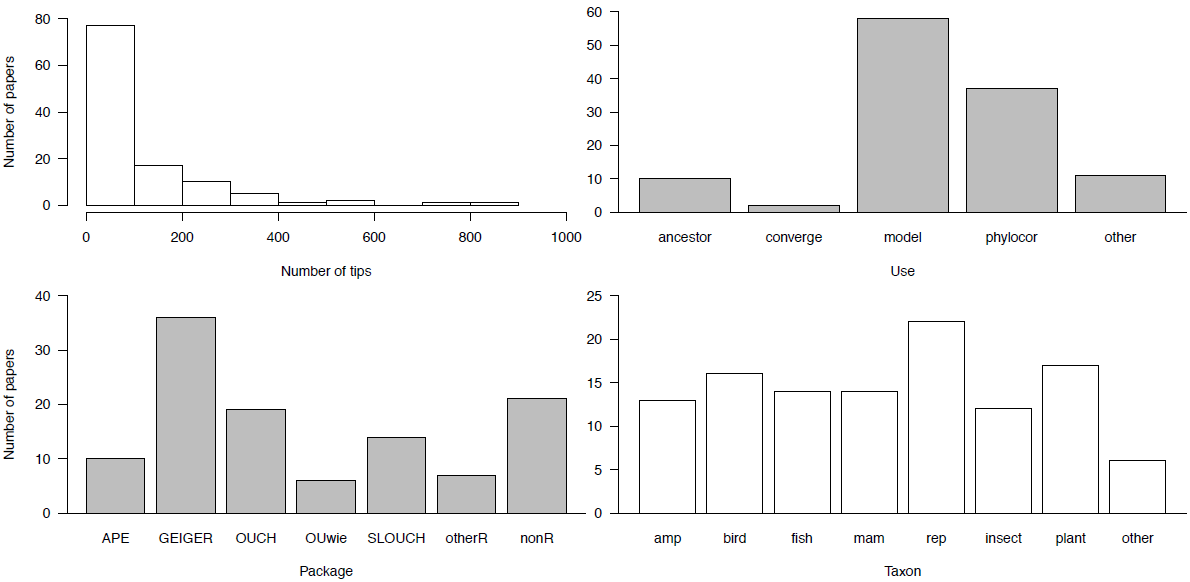
The median number of tips in phylogenies used (top left), the reason Ornstein Uhlenbeck (OU) models were fitted (top right), the R package used to fit the OU models (bottom left) and the taxon studied (bottom right) in each of the papers in the literature review (see (Table S1).

Use: ancestor = ancestral state reconstructions; converge = convergent evolution; model = model of evolution; phylocor = phylogenetic correction. ^†^Taxon: amp = amphibians; mam = mammals; rep = reptiles.

**Figure S2.**
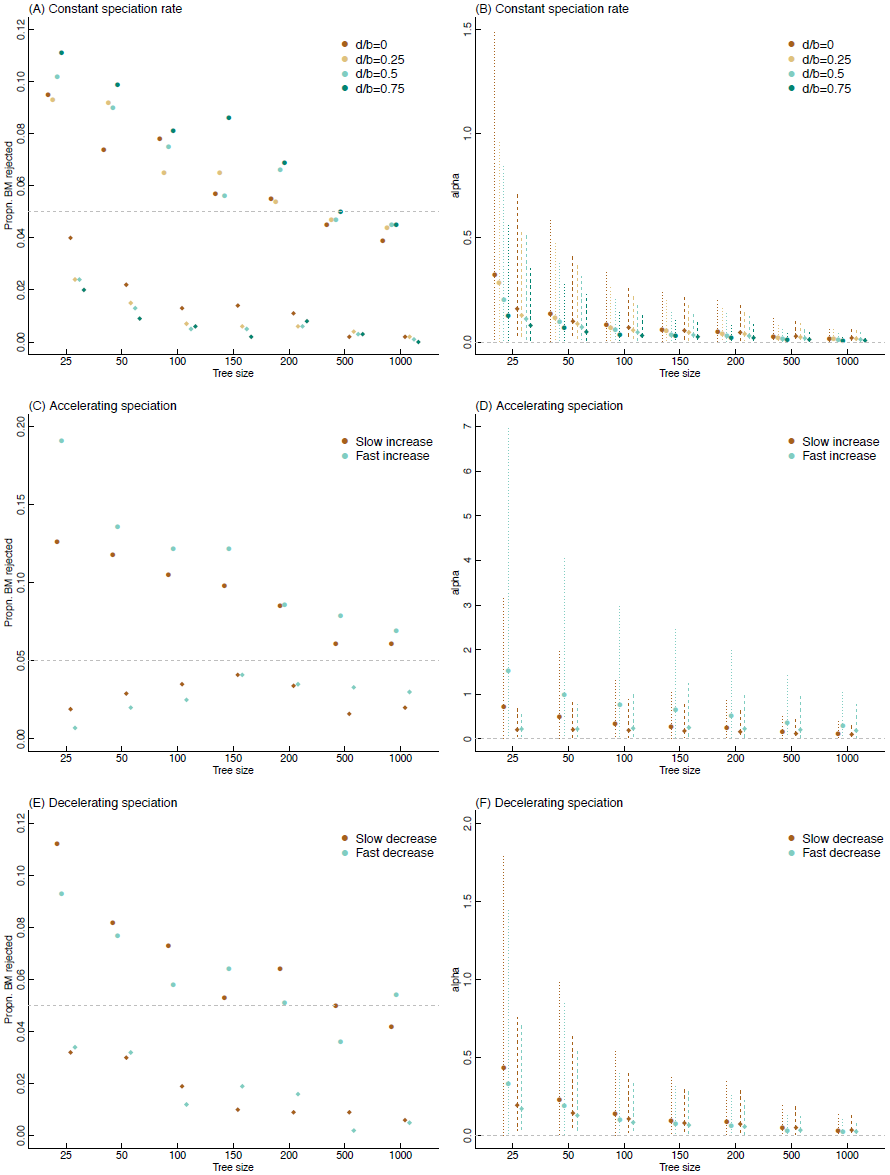
Rejection rates and estimation of *α* for different tree sizes and shapes. Rejection rates (a,c,e) for the OU model at multiple tree sizes for models fitted using ML (circles) and Bayesian (diamonds) methods. Estimates of *α* (b,d,f) based on ML (circles show means and dotted lines show 95% sampling interval from 1000 trees) and Bayesian (diamonds show means and dotted lines show 95% sampling interval of the modal value from the posterior distribution from 1000 trees). MCMC models fitted with an exponential prior (mean=1) on *α*.

**Figure S3.**
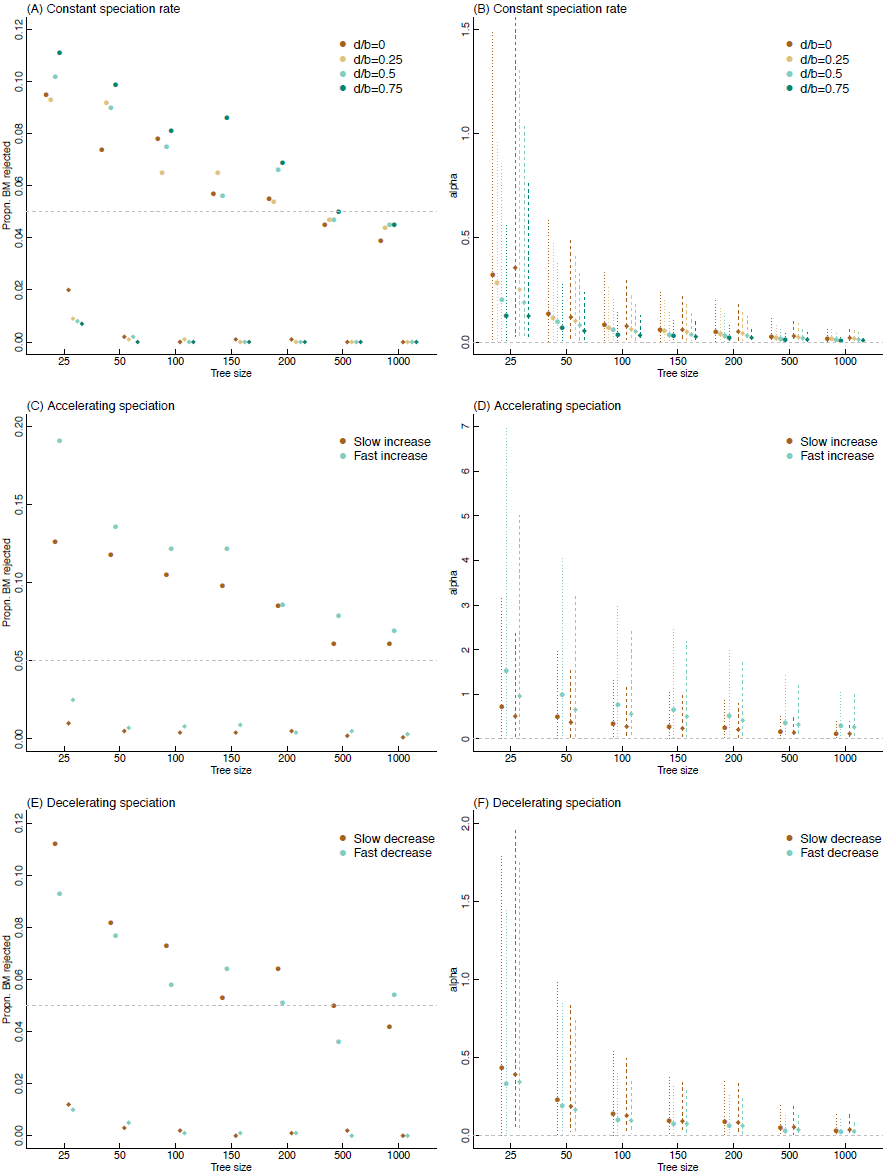
Rejection rates and estimation of *α* for different tree sizes and shapes. Rejection rates (a,c,e) for the OU model at multiple tree sizes for models fitted using ML (circles) and Bayesian (diamonds) methods. Estimates of *α* (b,d,f) based on ML (circles show means and dotted lines show 95% sampling interval from 1000 trees) and Bayesian (diamonds show means and dotted lines show 95% sampling interval of the modal value from the posterior distribution from 1000 trees). MCMC models fitted with uniform prior (0-20) on *α*.

**Figure S4.**
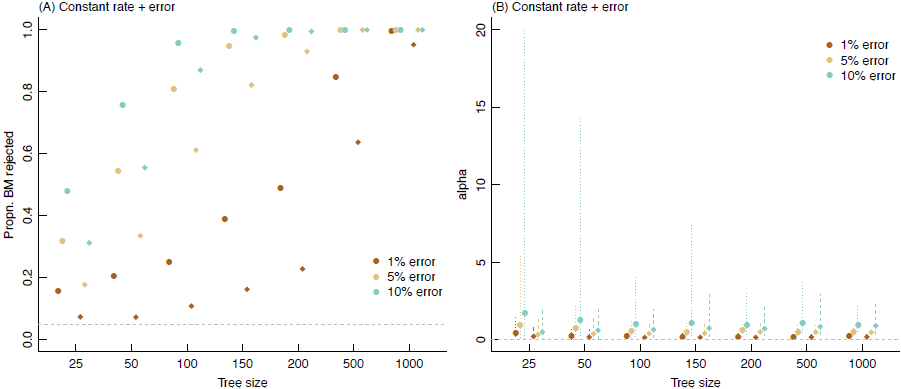
Rejection of BM with measurement error (a) and estimation of *α* (b). Symbols follow Figures S2 and S3. MCMC models fitted with an exponential prior (mean=1) on *α*.

**Figure S5.**
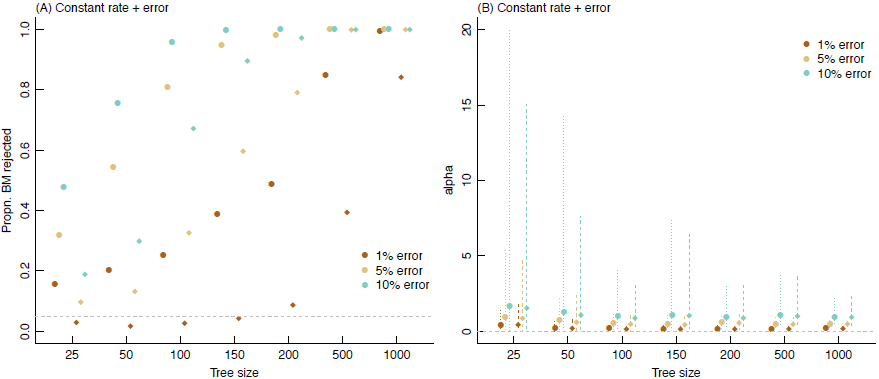
Rejection of BM with measurement error (a) and estimation of *α* (b). Symbols follow Figures S2 and S3. MCMC models fitted with uniform prior (0-20) on *α*.

**Figure S6.**
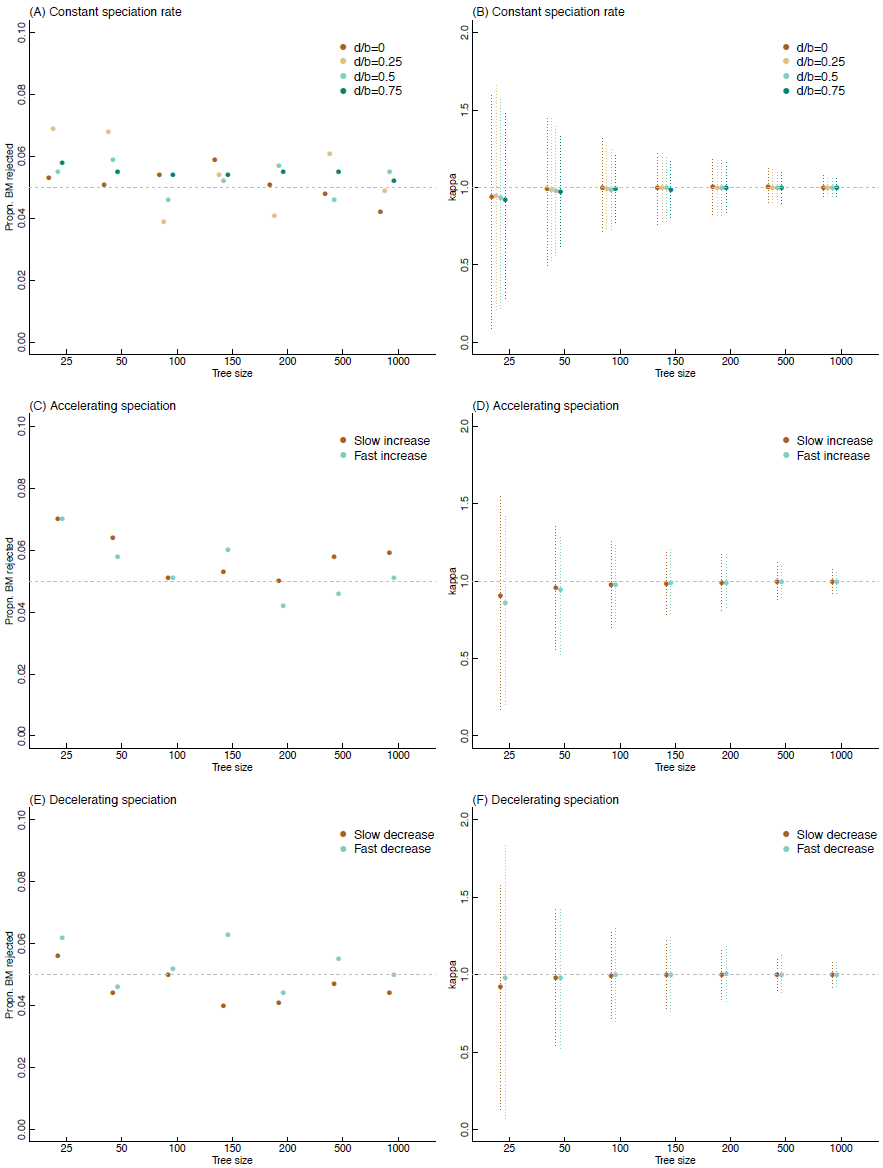
Rejection rates and estimation of Pagel's *κ* for different tree sizes and shapes. Rejection rates (a,c,e) for the *κ* model of speciational evolution at multiple tree sizes for models fitted using ML methods. Estimates of *κ* (b,d,f) based on ML with dotted lines showing 95% sampling interval from 1000 trees.

**Figure S7.**
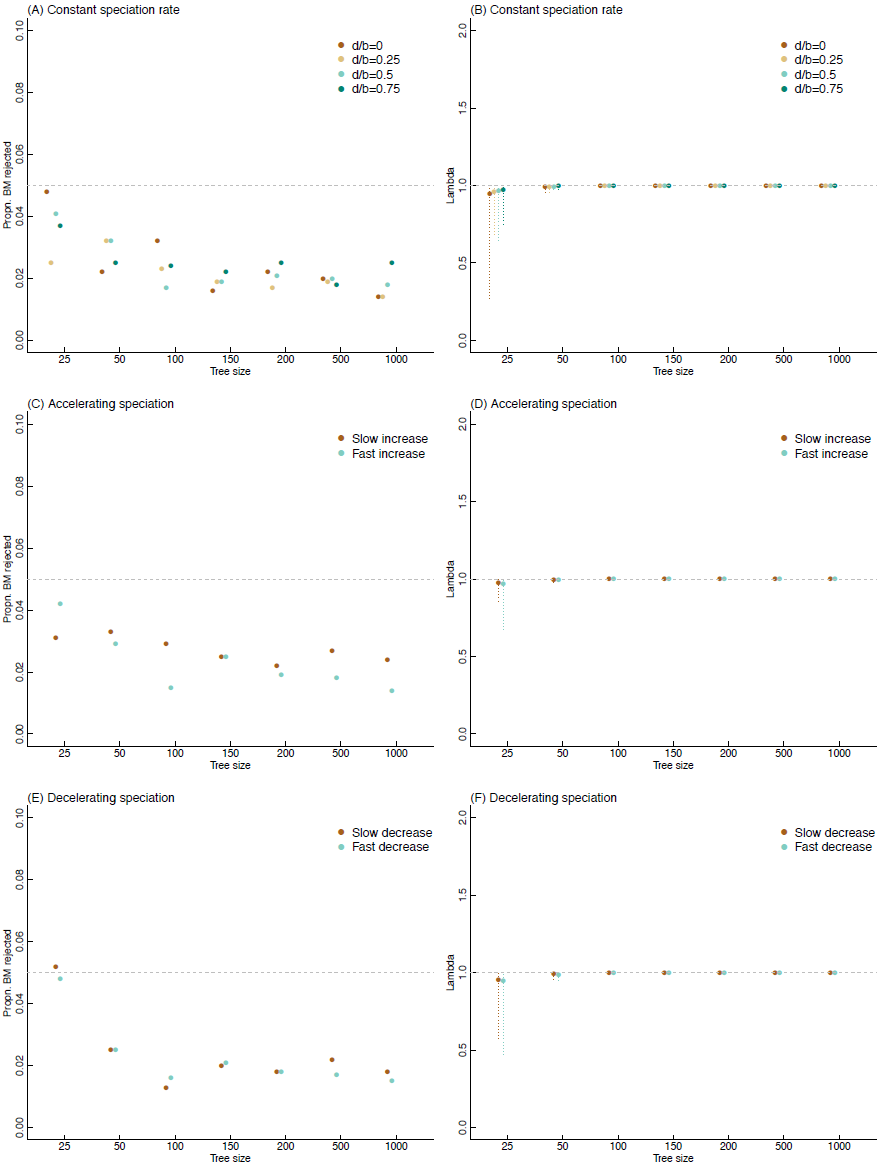
Rejection rates and estimation of Pagel's *λ* for different tree sizes and shapes. Rejection rates (a,c,e) for the *λ* model of phylogenetic signal at multiple tree sizes for models fitted using ML methods. Estimates of *λ* (b,d,f) based on ML with dotted lines showing 95% sampling interval from 1000 trees.

**Figure S8.**
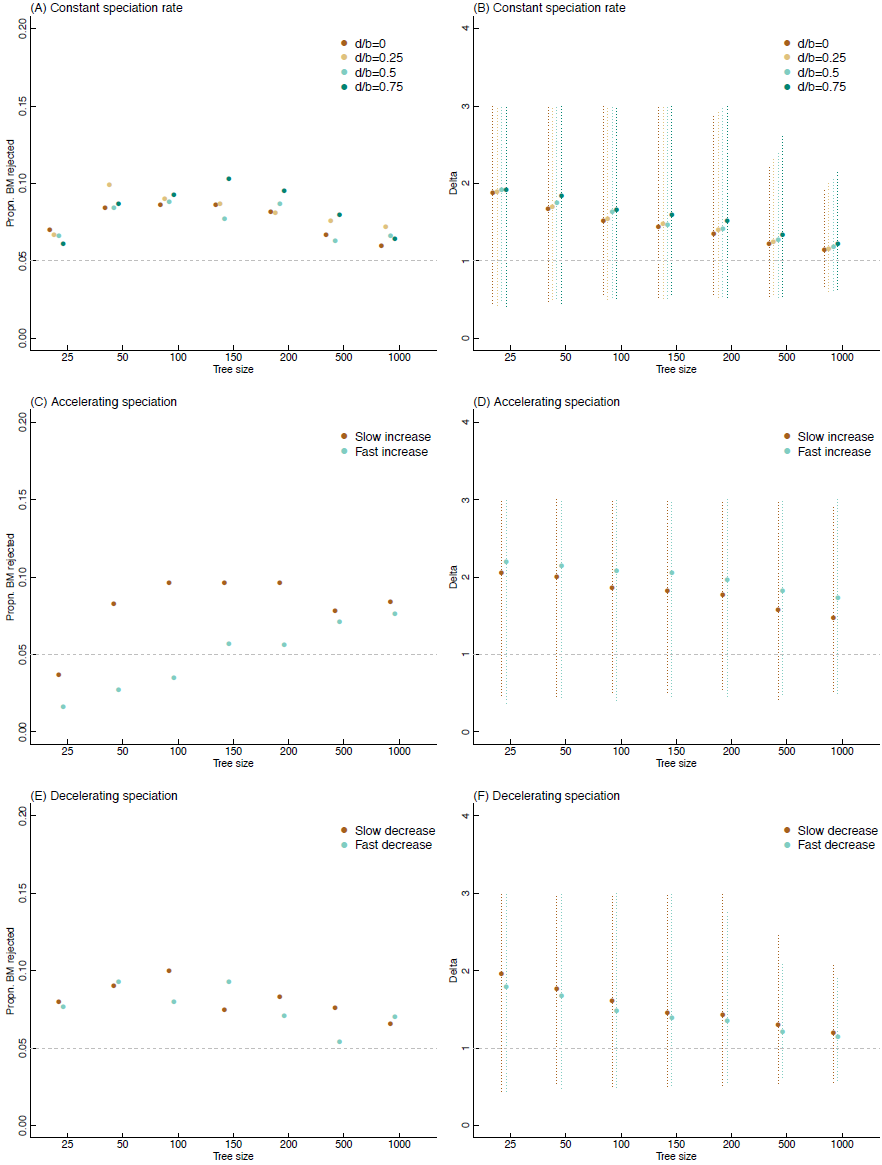
Rejection rates and estimation of Pagel's *δ* for different tree sizes and shapes. Rejection rates (a,c,e) for the *δ* model of accelerating/decelerating evolution at multiple tree sizes for models fitted using ML methods. Estimates of *δ* (b,d,f) based on ML with dotted lines showing 95% sampling interval from 1000 trees.

**Figure S9.**
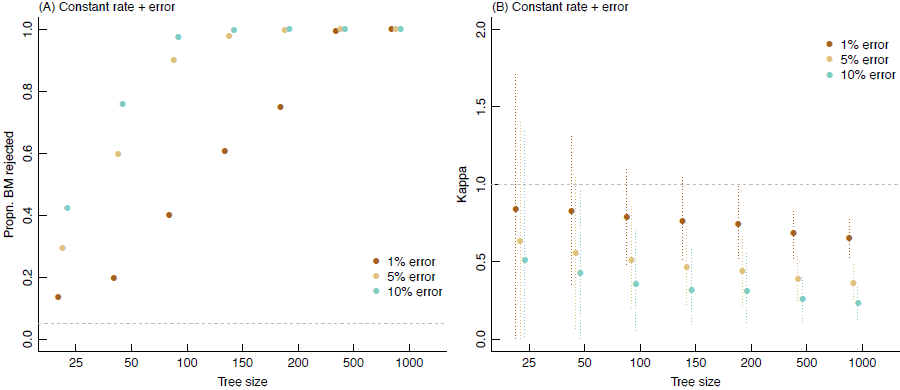
Rejection of BM with measurement error (a) and estimation of *κ* (b). Symbols follow Figures S6–S8.

**Figure S10.**
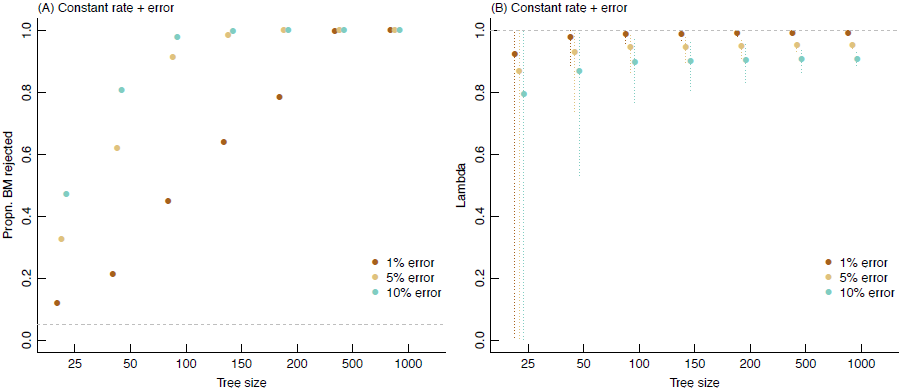
Rejection of BM with measurement error (a) and estimation of *λ* (b). Symbols follow Figures S6–S8.

**Figure S11.**
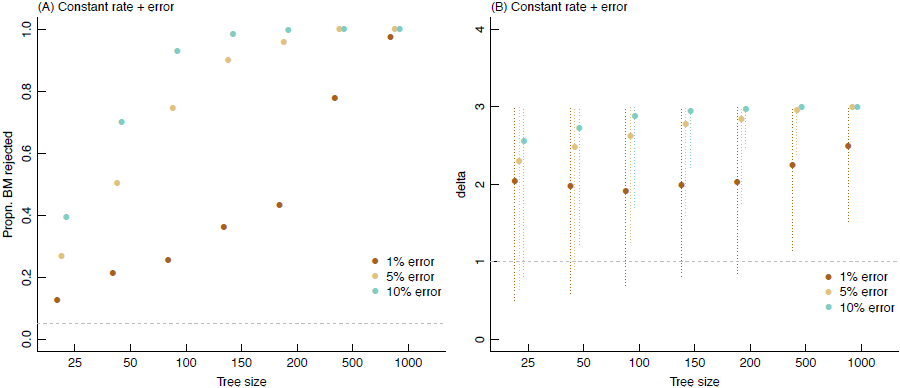
Rejection of BM with measurement error (a) and estimation of *δ* (b). Symbols follow Figures S6–S8.

